# Will they survive? Alarming circumstances of Asiatic cheetah (*Acinonyx jubatus venaticus*) in Iran’s drylands

**DOI:** 10.1101/2025.10.27.684750

**Authors:** Atie Taktehrani, Mahsa Shah Hosseini, Navid Gholikhani, Kaveh Hobeali, Mohammad Hossein Karimi, Negin Samadzadeh, Hamed Abolghasemi, Ali Ranjbaran, Ahmad Radman, Ahmad Safarzadeh, Morteza Pourmirzai, Mohammad S. Farhadinia

## Abstract

The Asiatic cheetah (*Acinonyx jubatus venaticus*), once widespread across West, South, and Central Asia, now survives only in Iran, where its population has declined to the brink of extinction. The current study synthesized 12 years (2012–2024) of monitoring data, including systematic, extensive camera trap surveys across 27 distinct sampling sessions in eight reserves (69,089 trap nights) supplemented by published records of cheetah occurrences on social media to assess the demographic and spatial patterns of this critically endangered subspecies. Our analysis indicates that a total of 24 adult Asiatic cheetahs were identified across the Northern and Southern Landscapes. However, no evidence of reproduction or new individual presence was obtained in the Southern Landscape for over a decade. Meanwhile, the Northern Landscape hosts the remaining population, likely fewer than 30 individuals. Between 2020 and 2024, at least 31 cubs were born in the northern population from six females. However, limited evidence of successful recruitment suggests minimal contribution to population recovery, as only 47.3% of monitored cubs survived beyond their first year. Asiatic cheetahs exhibit extensive mobility across the arid Northern Landscape, frequently traversing unprotected communal lands and a major highway, which increases their vulnerability. While camera trap data have proven effective for individual identification, they are limited in tracking fine-scale movement, emphasizing the urgent need for satellite telemetry. Interventions such as roadside fencing and wildlife underpasses along highways are essential to reduce mortality. These efforts should be complemented by broader conservation measures, including habitat protection and restoration, community-based management of unprotected lands, and enhanced anti-trafficking enforcement. Genetic concerns, especially low effective population size and inbreeding, pose additional threats to viability.

## Introduction

Conserving small populations of large carnivores presents considerable challenges due to compounded genetic, demographic, and ecological constraints. In small and isolated populations, inbreeding and genetic drift can reduce individual fitness, lower juvenile survival rates, and increase disease susceptibility (Hedrick & Fredrickson, 2010; Liberg et al., 2005). Such genetic erosion is often driven by founder effects and population bottlenecks, which limit adaptive potential and heighten extinction risk, particularly when numbers fall below minimum viable population thresholds (Marker et al., 2017; Benson et al., 2019).

These risks are especially pronounced in large carnivores such as cheetahs (*Acinonyx jubatus*), which naturally exist at low densities and require extensive, connected habitats for viable population function—factors that complicate reliable population estimation (Belbachir et al., 2015; Becker et al., 2017; Linden et al., 2020). Cheetahs are ecologically adapted to open landscapes such as savannas and arid plains, where they hunt fast-moving ungulates (Khalatbari et al., 2023; Weise et al., 2017). In arid regions, cheetahs cope with low prey densities by adopting nomadic movement patterns to exploit patchy resources; however, this increases their exposure to human-wildlife conflict, including persecution and road mortality (Farhadinia et al., 2016b). Their large home ranges, often exceeding 500 km^2^ in arid environments, combined with competition from larger predators, further threaten their survival (Marker et al., 2008b).

Male cheetahs, in particular, exhibit complex spatial behaviour tied to reproductive strategy (Melzheimer et al., 2018). Also, Melzheimer et al. (2020) reported that male cheetahs in Namibia begin as floaters with extensive home ranges (averaging 1,595 km^2^) that overlap with those of females (mean 650 km^2^) and smaller territories of resident males (mean 379 km^2^). Reproductive success in cheetahs is strongly limited by prey availability and habitat quality, with prolonged inter-birth intervals under resource scarcity constraining population growth (Marker et al., 2003; Wachter et al., 2011). Despite these ecological insights, population density and demographic structure remain poorly understood for many cheetah populations, particularly those inhabiting dryland ecosystems (Belbachir et al., 2015; Shams et al., 2025).

The Asiatic cheetah (*A. j. venaticus*), a critically endangered subspecies once widespread across West and Central Asia, has experienced a dramatic collapse since the 1970s due to habitat loss, prey depletion, and intensifying human-wildlife conflict (Hunter et al. 2007). Today, Iran remains its only known refuge (Farhadinia et al., 2016a). It is considered the rarest cheetah subspecies (Durant et al., 2017) and possibly one of the rarest felids globally. Since 2001, its total population has consistently been estimated at fewer than 70 individuals (Hunter et al., 2007; Farhadinia et al., 2017; Khalatbari et al., 2023).

Despite over two decades of extensive conservation efforts in Iran (Hunter et al., 2007; Farhadinia et al., 2017), the population shows no signs of recovery. Many historically occupied habitats have not yielded any confirmed cheetah detections in the past two decades, raising serious concerns about the subspecies’ long-term viability. This highlights the urgent need to understand the key demographic and spatial patterns shaping the survival of Asiatic cheetahs to guide current and future conservation strategies.

Since the start of Asiatic cheetah conservation efforts in 2001, we have provided decadal updates on both the species’ demographic (Farhadinia et al., 2016a, 2017) and spatial patterns (Farhadinia et al., 2016b), primarily using photographic evidence. Building upon these earlier assessments, we present an updated synthesis of the demographic status and spatial dynamics of the Asiatic cheetah, drawing on the most comprehensive dataset to date from 2012 to 2024. This long-term perspective offers critical insights into population persistence, distribution shifts, and the emerging challenges facing the species, informing future conservation priorities and recovery strategies.

## Materials and Methods

### Study Areas

In Iran, cheetahs are primarily distributed across low-canopy rangelands characterized by sparse vegetation (density <25%) and rugged terrains, reflecting a habitat preference shaped by topographic complexity. This contrasts with sub-Saharan African cheetahs, which typically inhabit flat to gently undulating grasslands, savannas, and shrublands, with only occasional presence in montane regions (Ahmadi et al., 2017). Based on inter-reserve movement data (Farhadinia et al., 2016b) and the spatial distribution of reserves, the Asiatic cheetah population is divided into two subpopulations located around the central deserts of Iran, the Northern and Southern landscapes. A third subpopulation, Kavir, has gone extinct very recently (Moqanaki & Cushman, 2016; Farhadinia et al., 2017). This study focuses on the Southern and Northern landscapes, particularly the latter, which currently represent the most critical habitats for the subspecies.

The Northern Landscape constitutes the primary breeding stronghold for Asiatic cheetahs in Iran. This landscape encompasses the Touran Biosphere Reserve (14,000 km^2^) and Miandasht Wildlife Refuge (840 km^2^), along with four adjacent protected areas (Fig. 1). These include Dorouneh Protected Area (667 km^2^), Khoshyeilagh Wildlife Refuge (1,380 km^2^), Chahshirin No-Hunting Area (680 km^2^), and Takhti Iran No-Hunting Area (350 km^2^). Among these, the Touran Biosphere Reserve and Miandasht Wildlife Refuge were the focal sites surveyed in this study.

**Fig. 1.**
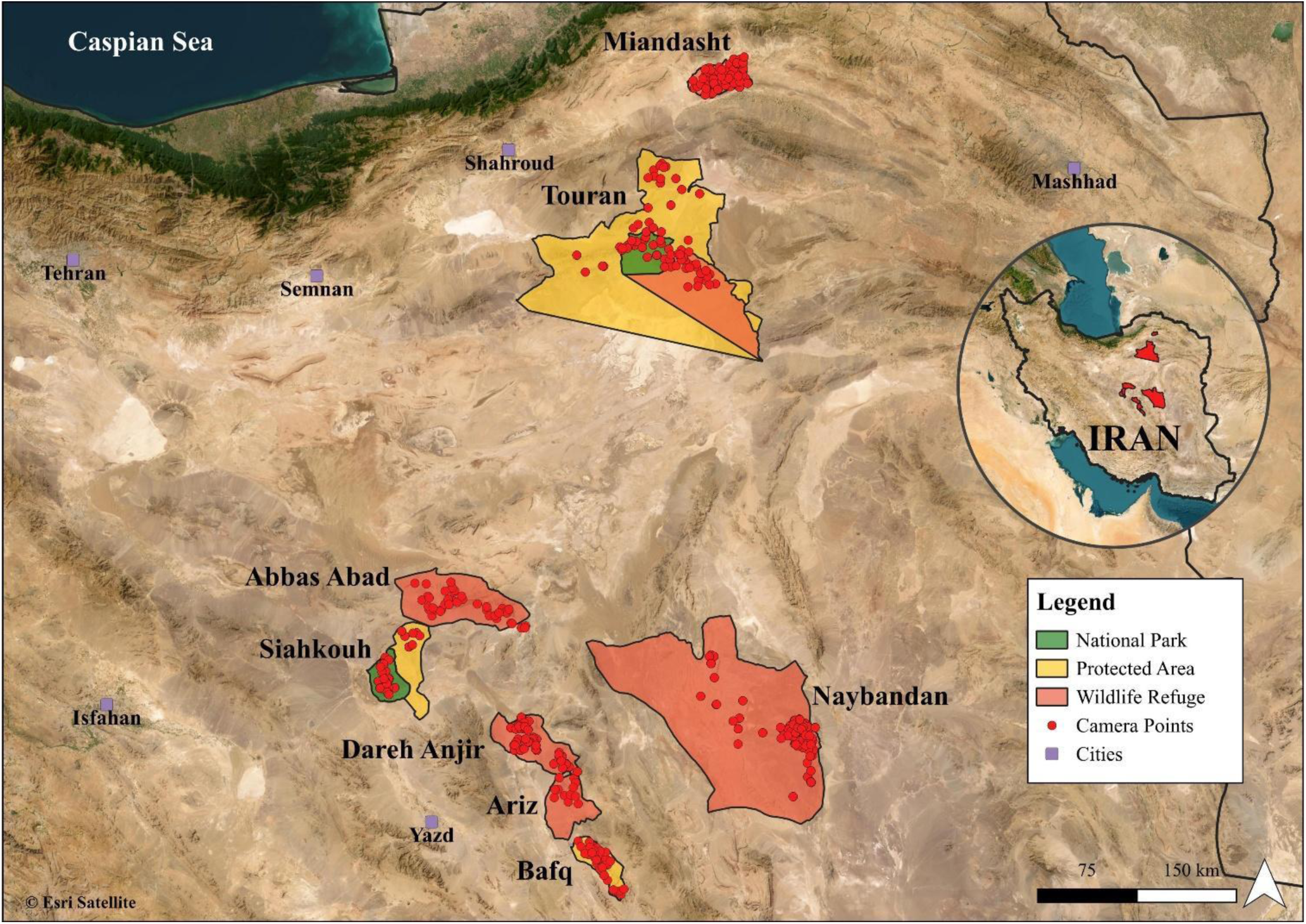
Map of the study area showing the spatial configuration of camera trap locations used for monitoring Asiatic cheetah (*Acinonyx jubatus venaticus*) populations in Iran. The map includes the Northern Landscape (Touran Biosphere Reserve and Miandasht Wildlife Refuge) and the Southern Landscape (Naybandan, Dareh Anjir, Ariz, and Abbas Abad Wildlife Refuges, as well as Siahkouh National Park, Siahkouh and Bafq Protected Area). Boundaries delineate the protected areas that were systematically monitored between 2012 and 2024.

The Southern Landscape encompasses 11 protected areas with confirmed cheetah presence. In this study, we surveyed six reserves, including Dareh Anjir Wildlife Refuge (1,753 km^2^), Siahkouh National Park and Protected Area (2,000 km^2^), Naybandan Wildlife Refuge (15,160 km^2^), Bafq Protected Area (885 km^2^), Abbas Abad Wildlife Refuge (3,050 km^2^), and Ariz Wildlife Refuge (960 km^2^) (Fig. 1).

### Sampling

Our synthesis is based on data from Systematic camera trap surveys (Table 1), supplemented with publicly available photographic records of Asiatic cheetahs by biologists, rangers, and citizen scientists published on social media between 2012 and 2024. These additional records were primarily captured by rangers patrolling and local community members who live around cheetah habitats. The systematic surveys covered eight reserves, deploying 692 camera trap stations for a total sampling effort of 69,089 trap-nights (Fig. 1).

**Table 1.**
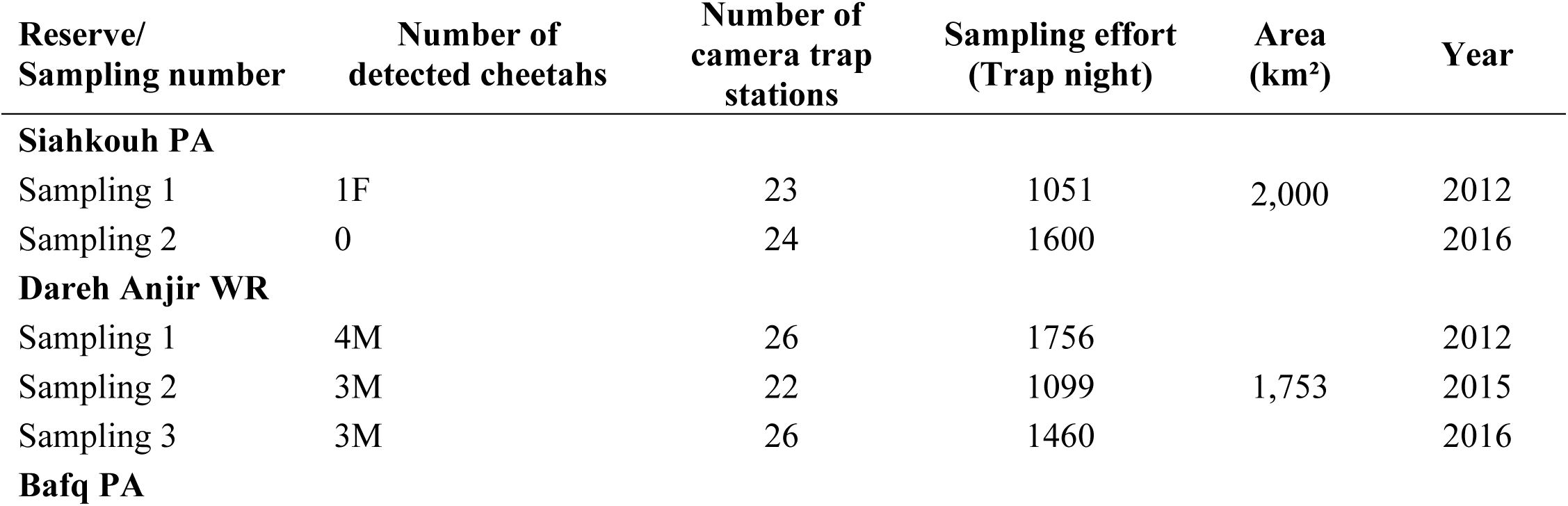
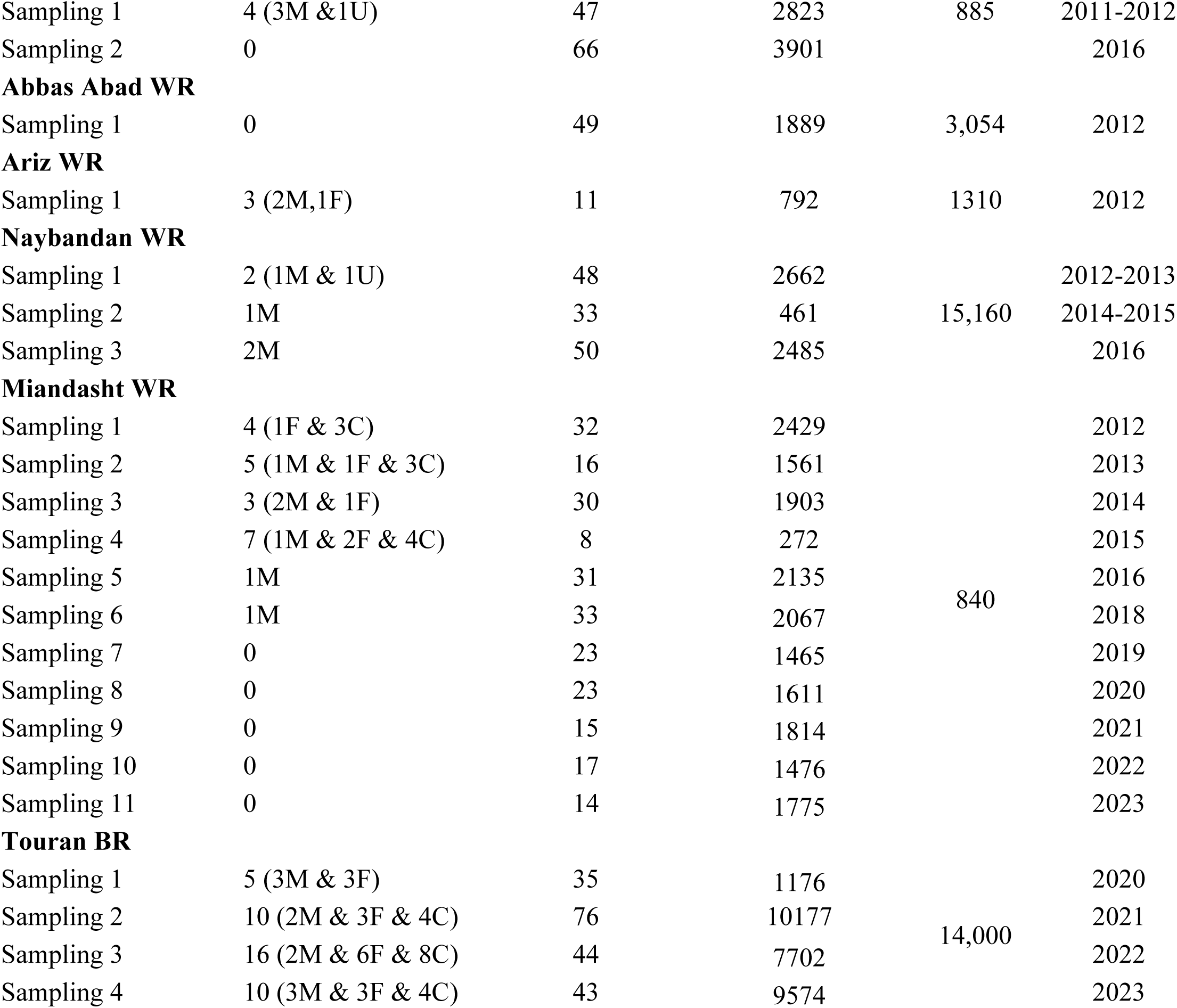
Details of individual Asiatic cheetahs confirmed in study areas between 2012 and 2023, including the monitoring season, the number of cheetahs (with sex and cub status), the number of camera traps, the number of trap nights, the reserve area, and the year. M=male, F=female, C=cub and U=Unknown sex. PA=Protected Area, WR=Wildlife Refuge, NP=National Park and BR=Biosphere Reserve.

Due to the extremely low cheetah population density, selecting optimal camera trap locations posed a significant challenge. Previous studies have shown that placing camera traps near male scent-marking sites (e.g., prominent trees or rocks) can lead to a sampling bias favouring male detections, as females tend to be more elusive (Farhadinia et al. 2017; Brassine and Parker, 2015; Shams et al., 2025). Therefore, to ensure more balanced sampling, camera traps were distributed across a range of habitat features, including mountain passes, flood paths, open plains, and other ecologically significant landscape features, guided by the local knowledge and expertise of reserve rangers. In drylands, water sources proved effective for camera deployment, particularly during the warmer months when female cheetahs, especially those with dependent cubs, show increased reliance on water (Farhadinia et al., 2023).

For individual identification, two camera traps were installed at frequently used crossing points, positioned on opposite sides of the trail to capture both flanks of passing cheetahs. Natural features such as rocks and bushes were used to funnel animals into the camera’s field of view, enhancing image clarity and identification accuracy. All individuals included in this study were identified based on their unique spot patterns. Individuals who could not be reliably identified were excluded from the analysis. Sex was determined visually, with males distinguished by the presence of external testicles.

We also developed a comprehensive photographic database of cheetah families. The number of cubs observed within the first six months after birth was used to estimate the average litter size per female. Cub survival metrics were calculated by continuously updating this database with camera trap records, supplemented by photographic records of Asiatic cheetahs published on social media, often taken by rangers or local community members. If a female was subsequently detected without her cubs within 17 months of the estimated birth date, a period typically associated with maternal care until independence (Laurenson, 1994), the litter was classified as lost. Litters that were not detected either during the first six months after birth or around the expected time of independence in the second year were excluded from survival rate analyses due to insufficient data. Mortality was recorded only when supported by photographic evidence confirming the death of a cheetah.

### Analysis

To track individuals over time, cheetah detections were compiled into a presence–absence matrix across survey years. Unless the cause of death was confirmed, all individuals were assumed to remain alive until the last year they were photographed. To summarize temporal persistence, the number of years each cheetah was detected was calculated, and the median and interquartile range (IQR) were used to report these values. This approach was adopted to reduce the influence of outliers, given the small sample size.

We then developed spatial movement patterns for each identified cheetah. Currently, no individuals have been detected in the Southern Landscape, and those previously present have already been characterized in earlier research (Farhadinia et al., 2016b). Consequently, this study focused exclusively on spatial patterns within the Northern Landscape, using camera trap data collected between 2020 and 2024, during which systematic monitoring was conducted.

Detection data were processed to identify unique individuals, and spatial analyses were conducted using QGIS version 3.40.4. Movement ranges were estimated using the Minimum Convex Polygon (MCP) method, which constructs the smallest convex polygon encompassing all recorded detection points for each individual. Only individuals with a minimum of 7 detections were included in the analysis.

We used annual counts of individual cheetahs detected from photographic evidence collected between 2014 and 2024 across the Northern Landscape. To model temporal trends in population metrics, we employed generalized additive models (GAMs) implemented in R (R Core Team, 2024) using the mgcv package (Wood, 2017). GAMs are flexible semiparametric regression models that accommodate non-linear relationships between predictors and response variables. GAMM captures non-linear relationships between variables and reports their complexity through effective degrees of freedom (EDF), where values close to 1 indicate linear effects, whereas higher values reflect nonlinear responses (Wood 2017). For each population variable, such as adult males, adult females, cubs of the year (<1 year), total adults, and mortality, we fitted a GAM including a smooth spline term with 8 basis functions to capture the potentially non-linear effect of year on demographic counts. The negative binomial family was used to account for overdispersed count data. Model predictions with 95% confidence intervals were generated for each year from 2014 to 2024. Model fit and assumptions were carefully evaluated to ensure adequate flexibility without overfitting.

## Results

### Demographic patterns

Over the 12-year study period, a total of 24 adult Asiatic cheetahs were identified in Iran (Fig. 2), comprising 14 males, 9 females, and one individual of unknown sex. 15 individuals were recorded in multiple protected and unprotected reserves, indicating a high degree of inter-reserve movement. The highest number of individuals was detected in Touran Biosphere Reserve, with 11 cheetahs (6 females and 5 males). Miandasht Wildlife Refuge accounted for five individuals, while eight cheetahs were recorded across the Yazd reserves (Dareh Anjir Wildlife Refuge, Siahkouh National Park, and Siahkouh Protected Area) and Naybandan Wildlife Refuge. Notably, sex ratios varied considerably across landscapes, ranging from a male-biased ratio in the Southern Landscape (6:1) to an equal sex ratio (8:8) in the Northern Landscape (Fig. 2), suggesting spatial differences in population structure and potential reproductive dynamics.

**Fig. 2.**
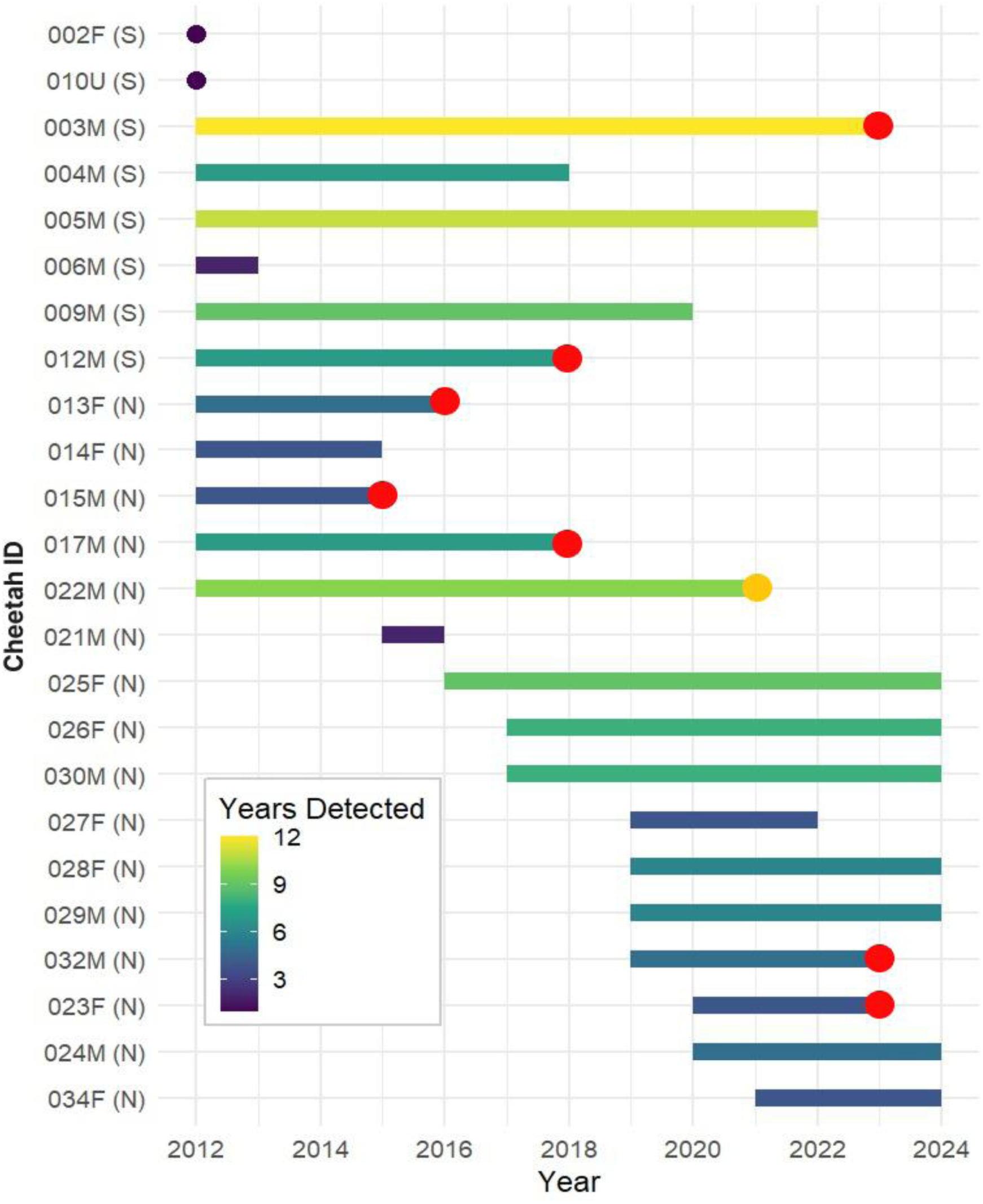
Abacus plot illustrating the annual detections of individuals of the Asiatic cheetahs in Iran (2012-2024). Each mark represents a confirmed sighting or detection event, highlighting temporal patterns and fluctuations in cheetah presence over the study period. The red dot indicates confirmed mortality, and the yellow dot represents that the individual was captured.

The median duration of camera-trap detections for individual cheetahs was 5.5 years (IQR = 4), with males showing a longer presence (Median = 7, IQR = 5) compared to females (Median = 4, IQR = 3). Since all individuals were recorded as adults, some likely reached the end of their natural lifespan during the study. Between 2020 and 2024, systematic monitoring in the Northern Landscape identified 11 cheetahs, of which 3 either died or were removed from the wild, leaving at least 8 adults in the population by the end of 2024. In the Southern Landscape, no new cheetah individuals have been recorded since 2011. The last known cheetah in the Southern Landscape was found dead in June 2023, and there have been no further detections since then (Fig. 2).

The last cheetahs known to be born in the Southern Landscape were a coalition of three brothers, 003M, 004M, and 005M, born in 2010. Equally important, the last known female in the Southern Landscape was recorded in 2012. In contrast, all documented breeding events of Asiatic cheetahs since 2010 have occurred exclusively in the Northern Landscape, where all known individuals have been repeatedly detected. However, no new adults have been identified in the past few years (Fig. 2).

Based on 14 identified families detected between 2012 and 2024 in the Northern Landscape, the average number of cubs per female was 2.7 ± SD 1.1 when they were first detected at the age of 3-6 months. In the Touran Biosphere Reserve, six females gave birth to 14 litters between 2020 and 2024, including three females that produced multiple litters. These litters resulted in a total of 31 cubs in the Northern Landscape, of which seven deaths have been confirmed (Table S1). Of the seven cheetah families we monitored using photographic records, six had at least one cub survive to its first year. However, only 47.3% (9 of 19 cubs) survived beyond their first year, with cubs that became independent from their mothers during this study period not included in this count.

Between 2012 and 2024, 21 cheetah mortalities were recorded, with all but two occurring in the Northern Landscape (Table S2). The leading cause was road collisions (8 deaths), followed by human-wildlife conflict (4 deaths). Other causes included orphaned cub removals (3), unknown factors (3), and single cases linked to illegal trade, veterinary malpractice, and natural causes.

The GAM revealed a significant temporal trend in the total number of detected cheetahs from 2014 to 2024 (edf = 1.47, χ^2^ = 11.75, p = 0.006), explaining 70.5% of deviance (adjusted R^2^ = 0.58) with a moderately non-linear pattern over time (Fig. 3). This indicates significant variation in detected cheetah numbers during the study period. In contrast, the smooth term for year was not significant for mortality (edf = 1.76, χ^2^ = 1.50, p = 0.46), with an intercept also non-significant (estimate = 0.20, p = 0.46). The mortality model explained only 23.2% of deviance (adjusted R^2^ = 0.11), suggesting limited predictive power. The small sample size (n = 11) may have reduced the ability to detect significant mortality trends (Fig. 3).

**Fig. 3.**
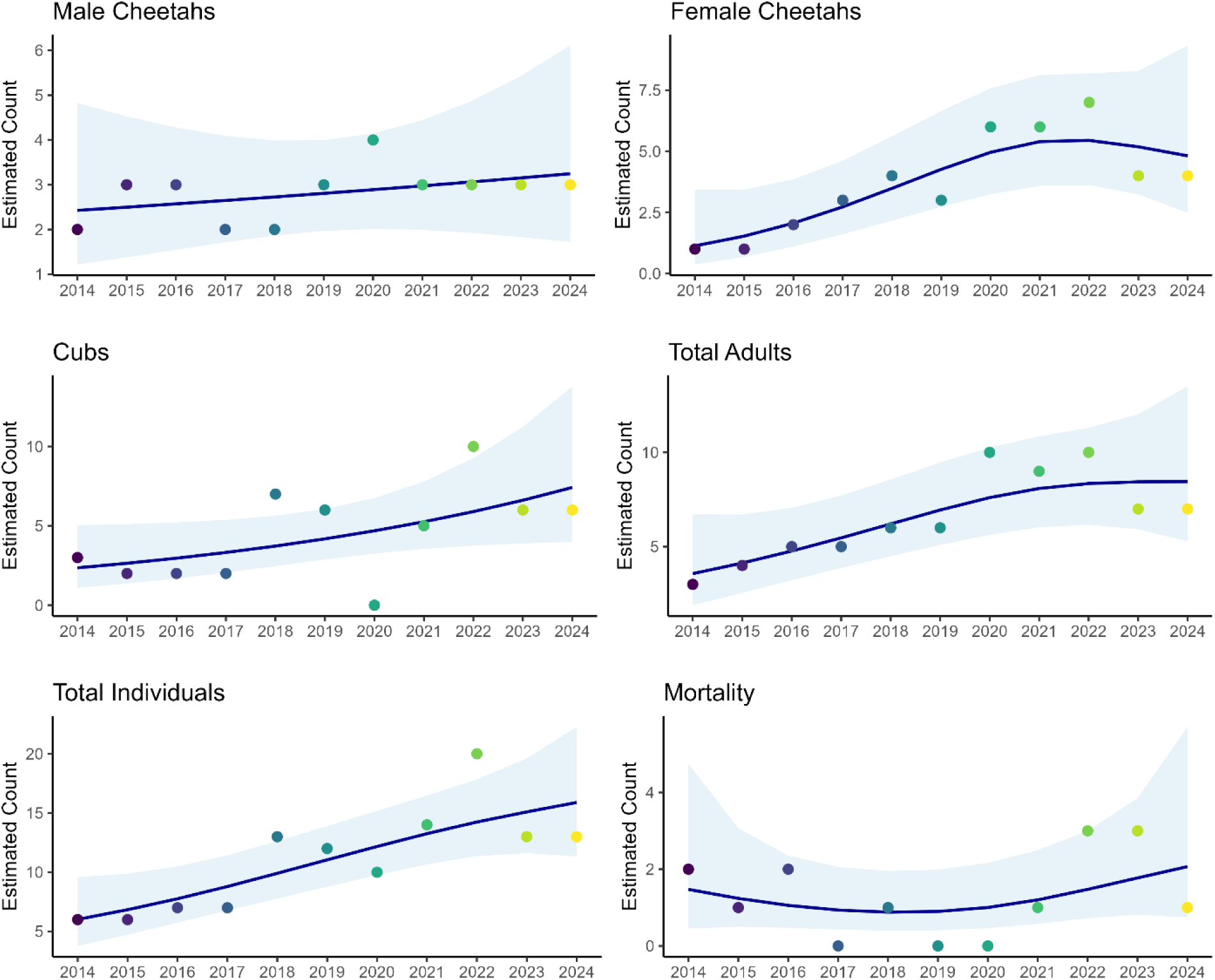
Temporal trends in Asiatic cheetah demographic metrics from 2014 to 2024 across the Northern Landscape in Iran. Each panel displays estimated counts for demographic parameters, including males, females, cubs, total adults, total individuals, and mortality, modelled using generalized additive models (GAMs) with a negative binomial error distribution. Solid lines show model-predicted trends, shaded ribbons represent 95% confidence intervals, and points indicate observed annual counts. Years are shown on the x-axis. Note that counts reflect detected individuals based on photographic evidence and may not represent the true population size.

### Spatial patterns

The median maximum distance between the farthest recorded locations for Asiatic cheetahs across the northern landscape was 81.9 km (IQR: 40.3 km), with males ranging farther than females (males: 83.1 km, IQR: 39.5 km; females: 70.6 km, IQR: 67.6 km). Similarly, the median MCP area for individual cheetahs was 1,354 km^2^ (IQR: 1,514 km^2^), with a larger range for males (1,524 km^2^, IQR: 2,021 km^2^) compared to females (1,184 km^2^, IQR: 1,354 km^2^). The spatial overlap of male and female cheetahs’ roaming areas, based on five years of camera trap data (2020–2024), is illustrated in Fig. 4

**Fig. 4.**
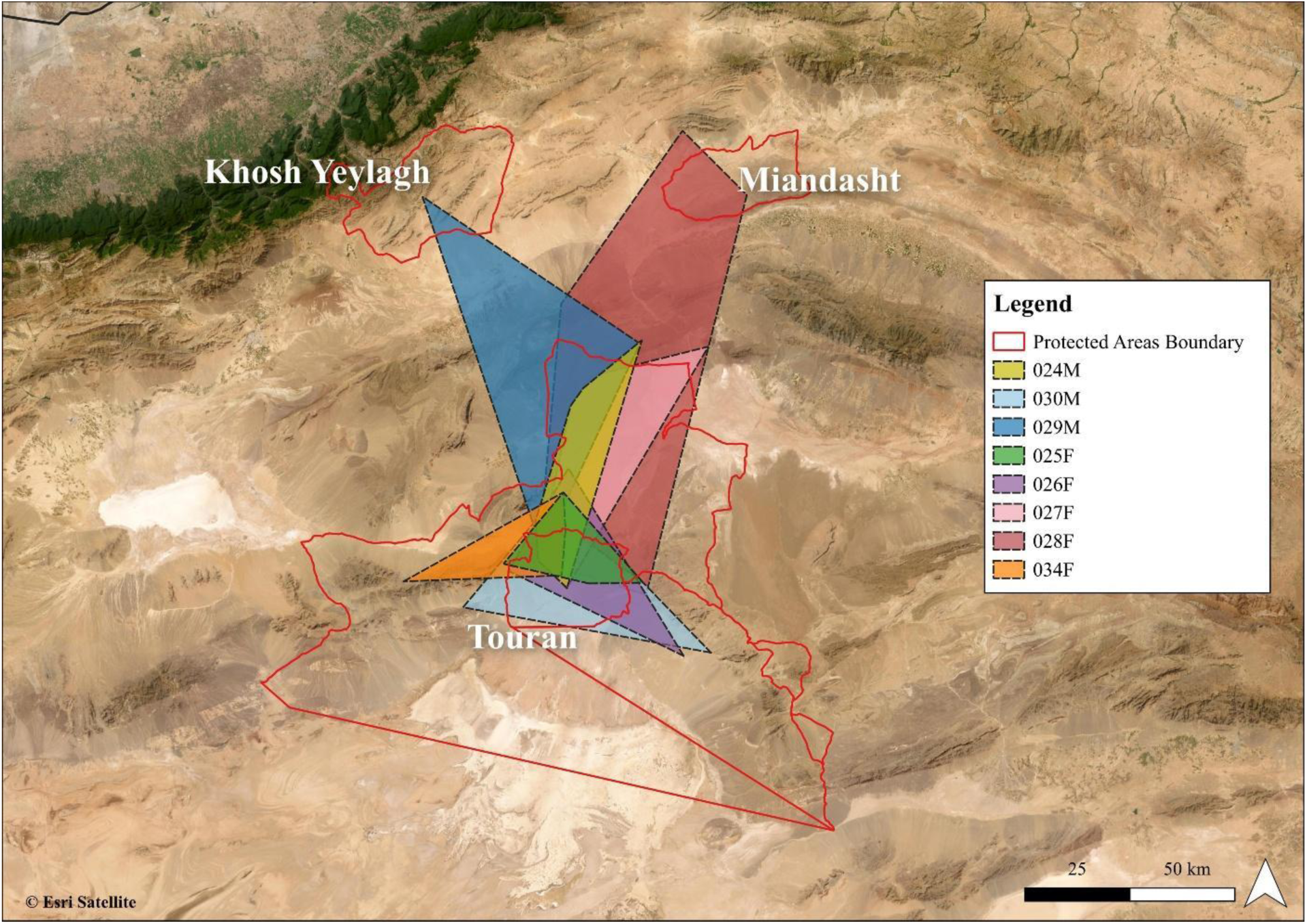
Ranging patterns of identified cheetahs in the Northern Landscape of Iran based on camera trap photos detected between 2020 and 2024. Each MCP colour refers to the relevant cheetah ID in Figure 2.

Of the eight identified Asiatic cheetahs, four individuals (025F, 026F, 030M, and 034F) were exclusively recorded within the Touran Biosphere Reserve. The remaining four (024M, 027F, 028F, and 029M) were initially detected in Touran but later observed in adjacent communal lands or other reserves (Table 2). Individual 027F, after separating dispersing from her mother and producing her first litter, was never seen again in Touran; all subsequent sightings occurred in unprotected communal lands. The location data we have for this female is from when she was within the Touran Biosphere Reserve. Thus, we have limited information on her movement range outside Touran.

Individual 028F displayed the widest range of movement: born in Touran, she gave birth to her first litter in communal lands, returned to Touran with her cubs, and later moved her second litter northward to Miandasht, ultimately covering 5,687 km^2^, the largest known range for any cheetah in Iran. Similarly, individual 029M, also born in Touran, went undetected for three years before being re-sighted in both the Touran Biosphere Reserve and the Khoshyeilagh Wildlife Refuge.

**Table 2.** Spatial patterns of Asiatic cheetah movement across the Northern Landscape based on camera trap detections between 2020 and 2024 in Iran.

| Cheetah ID | Max distance between farthest locations (km) | MCP (km<sup>2</sup>) | Reserves crossed |
| --- | --- | --- | --- |
| 024M | 83.1 | 1038 | Touran BR, communal land |
| 025F | 45.1 | 629 | Touran BR |
| 026F | 70.6 | 1184 | Touran BR |
| 027F | 89.7 | 1694 | Touran BR, communal lands |
| 028F | 149.3 | 5687 | Touran BR, communal land, Miandasht WR |
| 029M | 120.2 | 3059 | Touran BR, communal land, Khosh Yeilagh WR |
| 030M | 80.7 | 1524 | Touran BR |
| 034F | 58.6 | 686 | Touran BR |

## Discussion

Over the 12-year study period (2012–2024), only 24 adult Asiatic cheetahs were detected in Iran, despite extensive survey efforts totalling 69,089 trap nights across 27 distinct sampling sessions in eight reserves (Table 1). Notably, the confirmed mortality of seven individuals, accounting for 30.1% of known adults, was recorded by the end of 2024. Currently, there is no evidence of Asiatic cheetah presence in the Southern Landscape, with all recent detections limited to the Northern Landscape. Collectively, these findings highlight the extremely small and declining population of Asiatic cheetahs, casting serious doubt on the species’ long-term survival in the wild.

### Demographic patterns

Our data showed that the Northern Landscape, comprising Touran Biosphere Reserve and Miandasht Wildlife Refuge, remains the last breeding stronghold for the Asiatic cheetah. The high number of cubs born in the Northern Landscape between 2020 and 2024 (n=31) suggests a potentially high recovery capacity. However, a significant proportion of cubs were either confirmed dead (n=7, 22.5%) or remained unaccounted for (n=9, 29.0%). Given the lack of population recovery during this period, it is likely that many cubs did not survive to reach breeding age or contribute to population growth.

More critically, our findings revealed a worrying decline in cub survival rates compared to a decade ago in Iran’s Southern Landscape. Of the seven cheetah families we monitored using photographic records, six had at least one cub survive to its first year, like patterns observed in the 2010s, when at least one cub of all monitored families survived this milestone (Farhadinia et al., 2016a). However, overall cub survival has dropped sharply: just 47.3% (9 of 19 cubs) survived beyond their first year between 2020 and 2024 in the Northern Landscape, compared to 89.5% during the 2010s across the whole cheetah areas in Iran (Farhadinia et al., 2016a). For context, post-emergence cub survival in African populations varies widely, from 9.7% in the Serengeti (Laurenson, 1994) to 45.0% in the Kgalagadi Transfrontier Park (Mills & Mills, 2014), with some studies reporting survival until 14 months ranging from 54.5% to 95.8% (Laurenson, 1994; Mills & Mills, 2014).

The causes of mortality among Asiatic cheetah cubs remained poorly understood, while they seem to be randomly happening across the years. Confirmed cases include four cubs removed by humans, three trafficked for illegal wildlife trade, and one lost to apparent malnourishment. While African cub mortality is often driven by predation and starvation (Laurenson, 1994; Mills & Mills, 2014), anthropogenic threats appear to play a greater role in Iran’s remaining cheetah population. Mortality records from 2012 to 2024 (Table S2) show 21 documented deaths, 19 in the Northern Landscape and two in the south. Southern mortalities involved aging males, likely from natural causes, while most deaths affected females and cubs in the Northern Landscape, directly impacting reproductive potential. Historically (2001–2016), human-wildlife conflict was the leading cause of cheetah deaths, followed by road collisions and poaching (Farhadinia et al., 2016a, 2017). However, recent trends suggest a decline in conflict-related mortality, likely due to awareness and education campaigns, while road-related deaths have increased, possibly driven by expanding infrastructure (Mohammadi et al., 2023).

Although our GAM modelling indicated an apparent increase in the number of detected cheetah individuals over time, we consider this pattern more likely to reflect improved detection capacity rather than true population growth. Contributing factors may include increased ranger camera coverage, better knowledge of key cheetah locations, a greater sharing of photographic records by the public, and enhanced skills and equipment of our team.

Given the small sample size, we caution against drawing firm conclusions about population trends over the past 12 years. Nevertheless, the evidence suggests that a minimum level of breeding persists, with roughly three females producing litters each year (averaging 5–7 cubs). However, their contribution to long-term population recovery appears limited due to moderate-to-high cub mortality, much of which remains unexplained.

### Spatial patterns

Asiatic cheetahs demonstrated high spatial mobility across the drylands of the Northern Landscape, often traversing multiple reserves and unprotected communal lands. Of the eight adult cheetahs recently identified in Touran Biosphere Reserve, four moved into neighbouring landscapes. The median maximum distance between the farthest recorded detection points for individuals across the Northern Landscape was 81.9 km, with males generally traveling farther than females. The maximum recorded distance was 149.3 km for female individual 028F. In the Southern Landscape, multiple males were previously recorded at locations more than 150 km apart, with one male traveling up to 400 km between reserves (Farhadinia et al., 2016b). Similar long-distance movements have been observed in other arid regions: in Ahaggar Cultural Park, Algeria, the mean maximum distance travelled by two male cheetahs was approximately 45 km (Belbachir et al., 2015), while longer distances were also documented for males in the semi-arid farmlands of Namibia (Marker et al., 2008b; Melzheimer et al., 2020). The high mobility of Asiatic cheetahs and their large spatial requirement may also be a mechanism of inbreeding avoidance (Khalatbari et al. 2023).

Similarly, the median MCP area for individual cheetahs in the Northern Landscape was 1,354 km^2^, generally smaller than estimates from the 2010s, which were mostly recorded in the Southern Landscape (2,105.3 ± SE 778.6 km^2^; Farhadinia et al., 2016b). While both the current study and that of Farhadinia et al. (2016b) rely on camera trap data spread across expansive drylands, the estimated ranges remain considerably larger than those derived from the only GPS telemetry data available for Asiatic cheetahs (1,137 km^2^; Cheraghi et al., 2018). This highlights the species’ extensive spatial requirements and high mobility in arid environments. Comparable MCP estimates have been reported only in Namibia, where cheetahs in semi-arid farmlands occupy home ranges between 1,344 and 2,863 km^2^ (Marker et al., 2008b; Melzheimer et al., 2020). In contrast, cheetah ranges in sub-Saharan Africa are typically smaller (e.g., Houser et al., 2009), although Belbachir et al. (2015) reported a camera trap-based MCP estimate of 1,337 km^2^ for Saharan cheetahs, reinforcing the pattern of large home ranges in more arid ecosystems.

All cheetahs detected in this study were in the Touran Biosphere Reserve, underscoring its vital importance for the conservation of the species. However, individuals are frequently dispersed beyond the reserve boundaries, often crossing paved highways that fragment the landscape. These roads, while not impeding movement, significantly increase mortality risk, particularly from vehicle collisions, which have become a leading cause of death (Mohammadi et al., 2018; 2023). Notably, female 028 lost one of her male cubs in a road collision on the Tehran–Mashhad highway, one of Iran’s busiest transit corridors.

### Conclusion and management recommendations

Our analysis confirmed that the Asiatic cheetah population in Iran likely numbers fewer than 30 individuals, including juveniles, significantly lower than estimates from the past decade. This aligns with recent effective population size estimates (*N_e_* = 11–17; Khalatbari et al., 2023) and highlights an even more precarious conservation status than previously understood. At such critically low numbers, the population remains highly vulnerable to stochastic events, genetic deterioration, and demographic collapse. Based on our findings, we offer the following management recommendations:

1. The critically small population and signs of inbreeding threaten the Asiatic cheetah’s long-term viability (Khalatbari et al., 2023). Removing individuals before they breed, such as cubs or subadults, further reduces effective population size. Genetic rescue via admixture with African cheetahs has been recommended to counter inbreeding (Khalatbari et al., 2023) and should be seriously explored, starting with existing captive individuals to build a genetically robust backup population without further impact on wild cheetahs.
2. While camera traps offer valuable presence data, they fall short in revealing fine-scale movement patterns. Satellite telemetry is crucial to fill this gap, especially with fewer than 30 individuals remaining. GPS collars can provide essential insights into habitat use, movements, and survival, enabling more effective conservation strategies.
3. Road collisions, particularly along the Tehran–Mashhad highway, are now the leading cause of cheetah mortality, especially for mobile females and cubs. Urgent action is needed, including fencing, wildlife crossings, and improved road lighting, to reduce these fatalities (Mohammadi et al., 2018; 2023).
4. Illegal wildlife trade, driven by human–wildlife conflict and the removal of cheetah cubs from the wild, demands targeted interventions. These should include community education, strengthened enforcement against trafficking, improved livestock management, and enhanced habitat protection (Sardari et al., In press).
5. Cheetah movements increasingly occur outside formal reserves. In the Northern Landscape, unprotected transitional habitats have become vital for breeding and connectivity. Expanding community-based conservation into these areas is crucial for ensuring long-term population persistence (Ahmadi et al., 2020).

The collapse of southern populations illustrates the need to revise existing protocols to avoid repeating such losses in the north. While southern reserves should be maintained, conservation efforts must prioritise the Northern Landscape, with future reintroductions to the south considered only once population growth is achieved.

## Acknowledgement

We thank our partners, including the Iranian Department of the Environment, and our generous funders, especially Stichting SPOTS. We are also grateful to the many biologists, rangers, and citizen scientists who documented and publicly shared photographic evidence of Asiatic cheetahs on social media, which we accessed in accordance with applicable laws and ethical research standards. MSF was supported by Research England’s Expanding Excellence in England (E3) Fund and UK Research and Innovation. We also extend our appreciation to the rangers of the surveyed reserves for their invaluable field assistance and data collection.

## Supplementary Materials

**Table S1.** Cubs born, last recorded sightings, and known cub mortalities for Asiatic cheetah females in Miandasht Wildlife Refuge and Touran Biosphere Reserve, Iran (2012–2024).

| Reserve | Individual ID | Number of cubs | Birth year | Last seen with mother | Cub Mortalities |
| --- | --- | --- | --- | --- | --- |
| **Miandasht WR** | 013F | 3 | 2012 | Sep 2013 | 1 Cub died |
|  |  | 1 | 2015 | Sep 2015 | Unknown |
|  | 014F | 3 | 2015 | Oct 2015 | Unknown |
| **Total** | **-** | **7** | **-** | **-** | **1** |
| **Touran BR** | 025F | 3 | 2020 | May 2020 | Unknown |
|  |  | 2 | 2021 | Feb 2022 | 1 Cub died |
|  |  | 2 | 2024 | Unknown | Unknown |
|  | 026F | 3 | 2020 | Dec 2020 | 1 Cub died |
|  |  | 2 | 2021 | Aug 2021 | Unknown |
|  |  | 4 | 2023 | Unknown | 2 Cubs died |
|  | 027F | 4 | 2022 | Sep 2022 | Unknown |
|  | 028F | 4 | 2022 | June 2023 | 3 Cubs died |
|  |  | 2 | 2024 | Unknown | Unknown |
|  | 023F | 1 | 2022 | Nov 2022 | Unknown |
|  | 034F | 4 | 2024 | Unknown | Unknown |
| **Total** | - | **31** | - | - | **7** |

**Table S2.**
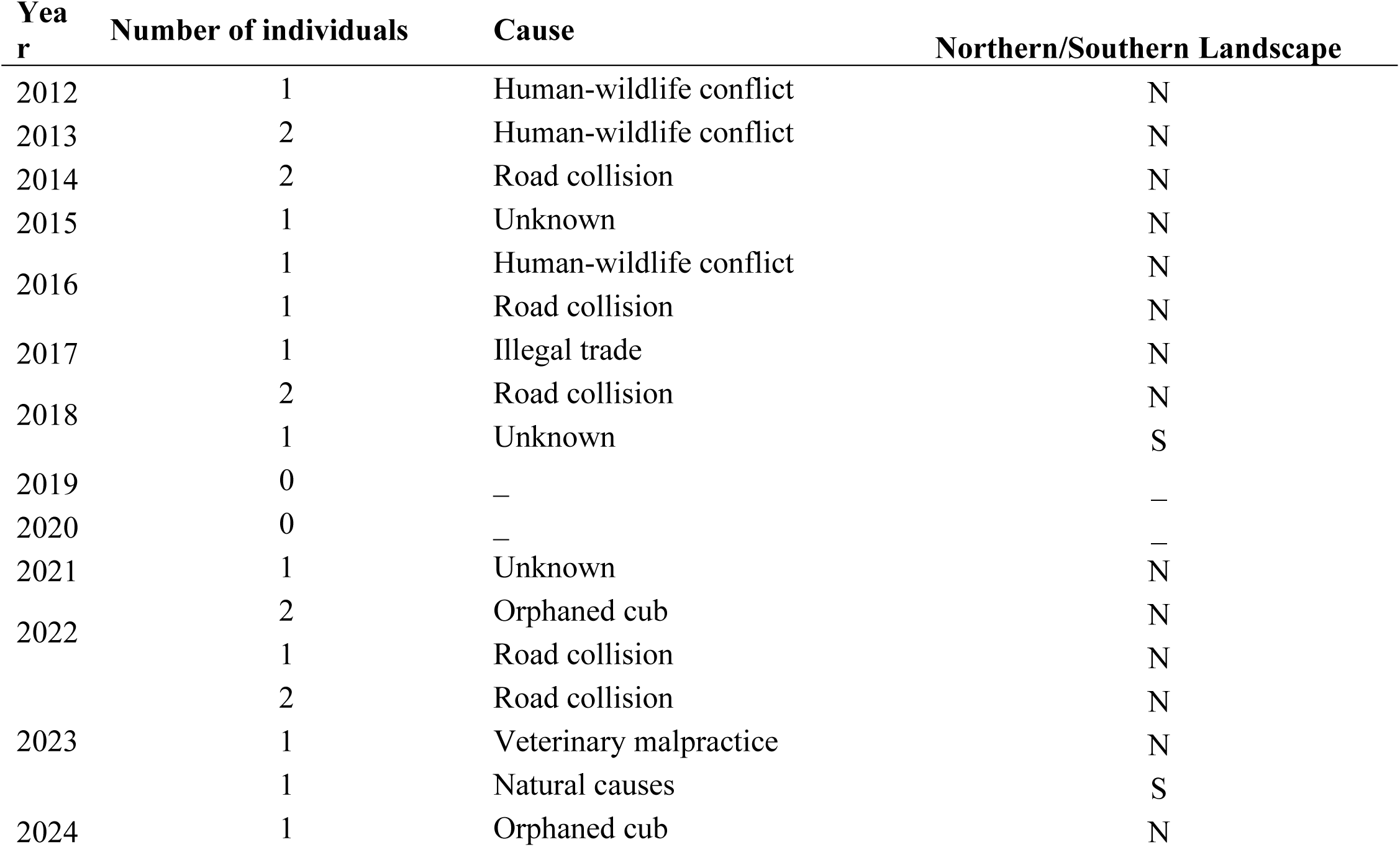
Details of cheetah mortalities in Iran between 2012 and 2024

## References

Ahmadi M, Nezami Balouchi B, Jowkar H, Hemami M-R, Fadakar D, Malakouti-Khah S, Ostrowski S. Combining landscape suitability and habitat connectivity to conserve the last surviving population of cheetah in Asia. Diversity Distrib. 2017;23(6):592–603. doi:10.1111/ddi.12560

Becker M, et al. Estimating past and present sex ratios of African cheetah populations. J Wildl Manage. 2017;81(2):234–245.

Belbachir F, Pettorelli N, Wacher T, Belbachir-Bazi A, Durant SM. Monitoring rarity: the critically endangered Saharan cheetah as a flagship species for a threatened ecosystem. PLoS One. 2015;10(1):e0115136.

Benson JF, Mahoney PJ, Vickers TW, Sikich JA, Beier P, Riley SPD, Ernest HB, Boyce WM. Extinction vortex dynamics of top predators isolated by urbanization. Ecol Appl. 2019;29(3):e01868.

Brassine E, Parker D. Trapping elusive cats: using intensive camera trapping to estimate the density of a rare African felid. PLoS One. 2015;10:e0142508.

Cheraghi F, Delavar MR, Amiraslani F, Alavipanah SK, Gurarie E, Fagan WF. Statistical analysis of Asiatic cheetah movement and its spatio-temporal drivers. Journal of Arid Environments. 2018 Apr 1;151:141–5.

Durant SM, Mitchell N, Groom R, Pettorelli N, Ipavec A, Jacobson A, et al. The global decline of cheetah and what it means for conservation. Proc Natl Acad Sci U S A. 2017;114:528–33.

Farhadinia MS, Akbari H, Eslami M, Adibi MA. A review of ecology and conservation status of Asiatic cheetah in Iran. Cat News Special Issue. 2016a;(10):18–26.

Farhadinia MS, Gholikhani N, Behnoud P, Hobeali K, Taktehrani A, Hosseini-Zavarei F, et al. Wandering the barren deserts of Iran: illuminating high mobility of the Asiatic cheetah with sparse data. J Arid Environ. 2016b;134:145–9.

Farhadinia MS, Hunter LTB, Jourabchian A, Hosseini-Zavarei F, Akbari H, Ziaie H, et al. The critically endangered Asiatic cheetah Acinonyx jubatus venaticus in Iran: a review of recent distribution, and conservation status. Biodivers Conserv. 2017.

Farhadinia MS, Nezami B, Ranjbaran A, Valdez R. Animal behavior informed by history: Was the Asiatic cheetah an obligate gazelle hunter? PLoS One. 2023;18(4):e0284593.

Hedrick PW, Fredrickson R. Genetic rescue guidelines with examples from Mexican wolves and Florida panthers. Conserv Genet. 2010;11(2):615–26.

Houser AM, Somers MJ, Boast LK. Home range use of free-ranging cheetah on farm and conservation land in Botswana. South African Journal of Wildlife Research-24-month delayed open access. 2009 Apr 1;39(1):11–22.

Hunter L, Jowkar H, Ziaie H, Schaller G, Balme G, Walzer C, et al. Conserving the Asiatic cheetah in Iran: launching the first radio-telemetry study. Cat News. 2007;(46):8–11.

Khalatbari L, Godinho R, Abolghasemi H, Hakimi E, Ghadirian T, Jowkar H, et al. The persistence of the critically endangered Asiatic cheetah relies upon urgent connectivity protection: a landscape genetics perspective. Conserv Genet. 2023;24(4):461–72.

Laurenson MK. High juvenile mortality in cheetahs (Acinonyx jubatus) and its consequences for maternal care. J Zool. 1994;234(3):387–408.

Liberg O, Andrén H, Pedersen HC, Sand H, Sejberg D, Wabakken P, et al. Severe inbreeding depression in a wild wolf population. Biol Lett. 2005;1(1):17– 20.

Linden DW, et al. Modeling cheetah population dynamics in southern Africa. Ecol Model. 2020;423:109013.

Marker L, Wilkerson AJP, Sarno RJ, Martenson JS, Breitenmoser-Würsten C, O’Brien SJ, et al. Molecular genetic insights on cheetah (Acinonyx jubatus) ecology and conservation in Namibia. J Hered. 2008a;99(1):2–13.

Marker LL, Dickman AJ, Mills MG, Jeo RM, Macdonald DW. Spatial ecology of cheetahs on north-central Namibian farmlands. J Zool. 2008b;274(3):226–38.

Marker L, Marnewick K, McNutt JW. Demography and population viability of cheetahs in small, isolated populations. Biodivers Conserv. 2017;26(8):1943–63.

Melzheimer J, Heinrich SK, Wasiolka B, Mueller R, Thalwitzer S, Palmegiani I, Weigold A, Portas R, Roeder R, Krofel M, Hofer H, Wachter B. Communication hubs of an asocial cat are the source of a human–carnivore conflict and key to its solution. Proc Natl Acad Sci U S A. 2020 Dec;117(33):202481117.

Melzheimer J, Streif S, Wasiolka B, Fischer M, Thalwitzer S, Heinrich SK, et al. Queuing, takeovers, and becoming a fat cat: Long-term data reveal two distinct male spatial tactics at different life-history stages in Namibian cheetahs. Ecosphere. 2018;9(6):e02308. doi: 10.1002/ecs2.2308

Mohammadi A, Almasieh K, Clevenger AP, Fatemizadeh F, Rezaei A, Jowkar H, et al. Road expansion: A challenge to conservation of mammals, with particular emphasis on the endangered Asiatic cheetah in Iran. J Nat Conserv. 2018;43:8– 18.

Mohammadi A, Ranjbaran A, Farhadinia MS, López-Bao JV, Clevenger AP. The Asiatic cheetah’s road to extinction. Science. 2023;382(6669):384.

Moqanaki EM, Cushman SA. All roads lead to Iran: Predicting landscape connectivity of the last stronghold for the critically endangered Asiatic cheetah. Anim Conserv. 2017;20(1):29–41.

QGIS Development Team. QGIS Geographic Information System. Version 3.40.4. Open Source Geospatial

R Core Team. (2024). R: A language and environment for statistical computing. R Foundation for Statistical Computing, Vienna, Austria. URL https://www.R-project.org/.

Shams A, Farhadinia MS, O’Riain MJ, Gaylard A, Smit M, Fraticelli C, et al. Perilous state of critically endangered Northwest African cheetah (Acinonyx jubatus hecki) across the Sudano-Sahel. Anim Conserv. 2025;28(2):208–23.

Wachter B, et al. Reproductive history and absence of predators are important determinants of reproductive fitness: the cheetah controversy revisited. Conserv Lett. 2011;4(1):47–54.

Weise FJ, et al. The distribution and numbers of cheetah (Acinonyx jubatus) in southern Africa. PeerJ. 2017;5:e4096.

Wood, S.N. (2017) Generalized additive models: an introduction with R. Chapman and hall/CRC. Available at: https://www.taylorfrancis.com/books/mono/10.1201/9781315370279/generalized-additive-models-simon-wood (Accessed: 3 August 2025).

